# Frequency-specific transcranial neuromodulation of oscillatory alpha power alters and predicts human visuospatial attention performance

**DOI:** 10.1101/2020.08.04.236109

**Authors:** S. K. Kemmerer, A.T. Sack, T.A. de Graaf, S. ten Oever, P. De Weerd, T. Schuhmann

## Abstract

Unilateral transcranial alternating current stimulation (tACS) at alpha frequency modulates the locus of spatial attention. However, the neural mechanisms by which tACS influences spatial attention remain poorly understood. Here, we applied high-definition tACS at the individual alpha frequency (IAF), two control frequencies (IAF+/-2Hz) and sham to the left posterior parietal cortex and measured its effects on visuospatial attention performance as well as alpha power (using electroencephalography, EEG). Our results revealed a leftward lateralization of alpha power relative to sham. At a high value of leftward alpha lateralization, we also observed a leftward attention bias, which differed from sham. Moreover, the magnitude of the alpha lateralization effect predicted the attention bias. These effects occurred for tACS at IAF but not for the control frequencies. This suggests that tACS operates through oscillatory interactions with ongoing brain rhythms in line with the synchronization theory. Our results also highlight the importance of personalized stimulation protocols, especially in potential clinical settings.

## Introduction

As the number of visual stimuli in the visual world exceeds the processing capacity of our brain, we have to filter the visual input. *Visuospatial attention* - a form of visual attention - helps us to select stimuli for enhanced processing based on their location in space [1–4]. It thereby acts as an attentional filter enabling us to prioritize some stimuli over others. A visuospatial attention *bias* describes the tendency to pay more attention towards one side in space compared to the other side. We all display a visuospatial attention bias, which is so small that it can only be revealed in experimental settings [5]. A more severe and pathological visuospatial attention bias, called visual hemineglect, can emerge after unilateral damage to the attention network due to stroke or traumatic brain injury [6,7]. Although basic visual and motoric functions may be completely intact, hemineglect patients have difficulty perceiving and responding to stimuli on the contralesional side [8,9].

On a neural level, natural as well as pathological visuospatial attention biases are associated with an interhemispheric asymmetry in oscillatory alpha (7-13Hz) power over posterior sites [10,11]. Similarly, dynamical shifts of visuospatial attention to either hemifield lead to a lateralization of occipitoparietal alpha power with higher alpha power ipsilateral to the attentional locus [12–15]. In this context, it has been postulated that alpha oscillations could serve as an attentional inhibition mechanism, enabling the selective processing of relevant stimuli by suppressing distracting incoming sensory information [16–18]. However, based on correlational electroencephalography (EEG) data it is difficult to draw definitive conclusions about the functional relevance of alpha oscillations. It remains possible that alpha oscillations are an epiphenomenon; a by-product of another attentional mechanism. To demonstrate a direct relationship between alpha oscillations and visuospatial attention, it is necessary to modulate alpha power and show that this leads to a change in visuospatial attention performance.

Transcranial alternating current stimulation (tACS) is a non-invasive brain stimulation technique, which uses alternating electrical currents to increase the power of brain oscillations [19–21]. Numerous studies have reported effects of tACS on perception [22–24], cognitive functions [26–28] as well as motor control and learning [29–38] (for a recent review see [39]). Furthermore, experiments that combine tACS with EEG indicate that alpha power over both hemispheres can be enhanced through medial occipitoparietal tACS at alpha frequency [40–42]. Building upon these findings, we recently applied high-density (HD) tACS at 10Hz to the left posterior parietal cortex (PPC) with the aim of modulating the visuospatial attention locus. In line with our hypothesis, we demonstrated that tACS at 10Hz induces a visuospatial attentional leftward bias relative to sham [43]. Even more recently, this effect was replicated and extended by Kasten and colleagues [44] who showed that the effect of tACS at alpha frequency on the visuospatial attentional locus is inverse to the effect of tACS at gamma frequency and can only be found during left but not right hemispheric stimulation. Similar effects were report in the auditory domain by Wöstmann and colleagues [45] as well as Deng, Reinhart and Choi [46] who found an ipsilateral shift of auditory spatial attention during unilateral tACS at 10Hz. This suggests that alpha oscillations indeed play a causal role in attentional control [14–16,18,47,48]. The results also indicate a potential clinical application of tACS, in which unilateral tACS at alpha frequency is used to correct the pathological attention bias of hemineglect patients.

While these tACS studies on spatial attention show a clear behavioral stimulation effect, they did not include neuroimaging to verify the underlying neural effects. Spatial attention tasks can reveal information about the behavioural tACS effect, but EEG or magnetoencephalography (MEG) measurements in the same paradigm are necessary to confirm that the power of the targeted oscillation was indeed enhanced as intended. Without such measurements, it is impossible to conclude with certainty that the behavioural tACS effects were driven by the assumed changes in oscillatory power. It is therefore still unclear whether the tACS effects on spatial attention are driven by modulations of alpha power or other neural changes, especially because tACS-induced alpha power enhancements have only been demonstrated with central but not lateralized tACS montages. Furthermore, the extent of the stimulation frequency-specificity of tACS has not yet been fully explored. The theoretical framework of synchronization [49] predicts that rhythmic stimuli preferentially enhance an oscillation if they are applied at the intrinsic dominant frequency. With increasing deviation of the stimulation frequency from the intrinsic dominant frequency, the stimulation effect is expected to diminish and to approach zero. In line with these ideas, it has been proposed that tACS operates via synchronization of neural oscillations to the alternating current [21]. However, we are not aware of any tACS experiment that compared the effects of stimulation at the intrinsic dominant frequency to stimulation at close flanking control frequencies. A frequency specific stimulation effect, limited to tACS at the intrinsic dominant frequency but not close flanking frequencies, would therefore provide important support for the synchronization theory [49] as the neural basis for tACS effects.

Here, we tested the effect of left parietal HD-tACS at alpha frequency on visuospatial attention performance and oscillatory alpha power in participants of various age groups spanning from adolescence to mature adulthood. We applied tACS at the IAF, two control frequencies IAF+/-2Hz as well as sham (placebo) stimulation and each participant underwent all four stimulation conditions (IAF, IAF+2Hz, IAF-2Hz, sham) in separate sessions and randomized order (Fig 1A). We used a high-definition (HD) ring electrode montage, which creates a focused electrical field and thereby allows for precise targeting of the left posterior parietal cortex (Fig 1B). During stimulation, we assessed visuospatial attention performance with a spatial cueing (Fig 1C) and a detection task (Fig 1D). Before and immediately after stimulation, we collected resting state electroencephalography (EEG) data to measure the effect of tACS on offline alpha power (see methods section for details).

**Fig 1.**
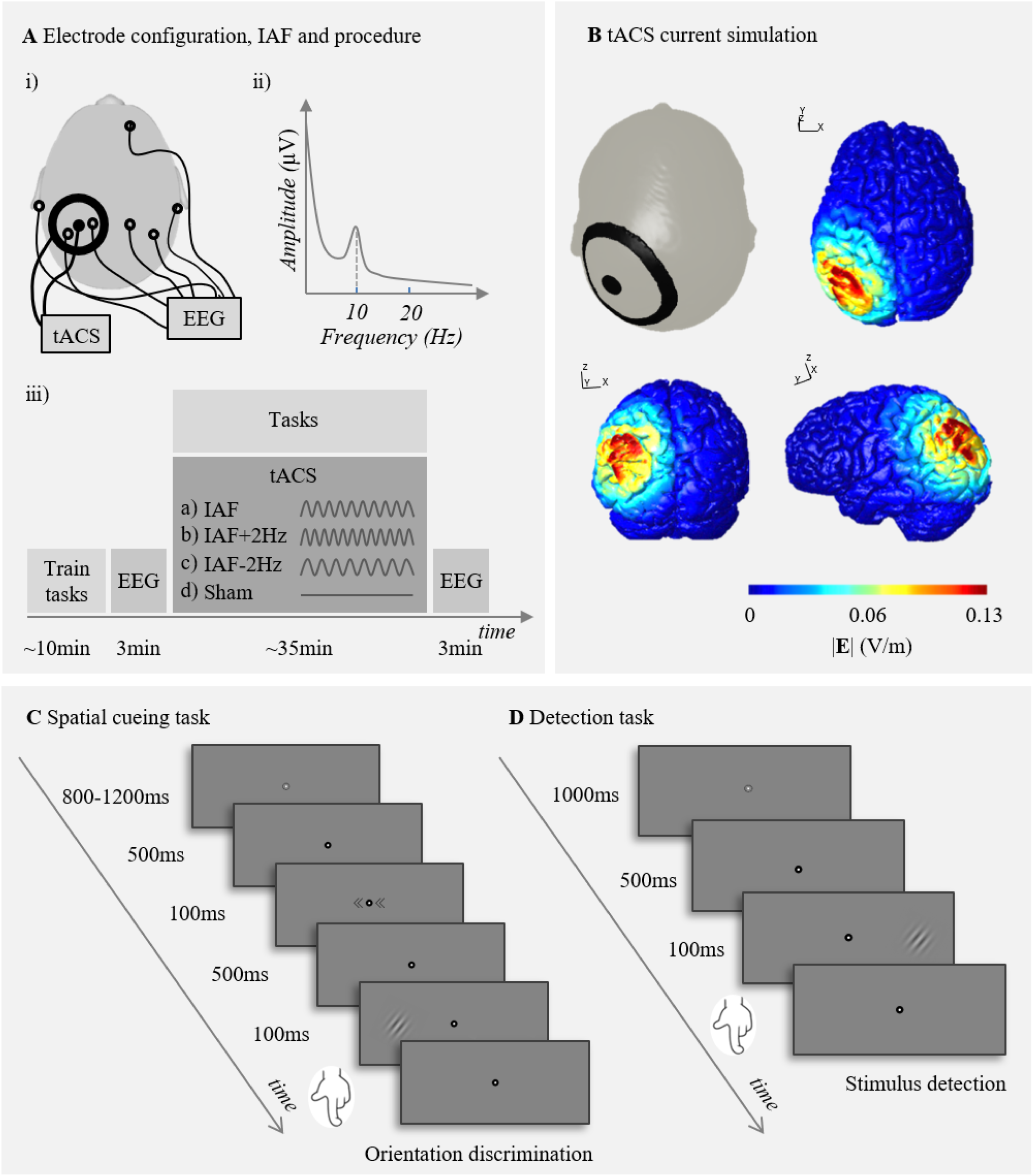
Electrode configuration, IAF, procedure, current simulation and example trials of the attention tasks. (A, i) *tACS and EEG electrode configuration*. The small tACS disk electrode was placed on the left posterior parietal cortex (P3) and the large ring electrode was centered around it. In between the disk and the ring electrode, we mounted two single EEG electrodes and mirrored to those also two EEG electrodes in the right hemisphere. (A, ii) *IAF*. This exemplary EEG power spectrum depicts power over frequency (Hz). The dashed line marks the alpha peak, also called individual alpha frequency (IAF). (A, iii) *Procedure*. At the beginning of each session, participants completed a shortened practice version of the attention tasks. Then, three minutes of resting state EEG data were measured while the participants kept their eyes closed. This was followed by tACS at (a) IAF (b) IAF+2Hz (c) IAF-2Hz or (d) sham, while the participants performed both attention tasks. Each participant underwent all four stimulation conditions in separate sessions. After completion of both tasks, the tACS device was switched off and three minutes of resting state EEG data were measured again. (B) *Current simulation results*. The norm electric field (V/m) is depicted on an example brain from a transverse, coronal and left sagittal view. (C) *Example trial of the spatial cueing task*. A given trial started with a fixation period during which only a bullseye fixation point was shown. Then, arrow heads pointing to the left, right or both sides, were presented. These symbolic cues flanked the central fixation point and were used to evoke endogenous shifts of visuospatial attention. After another brief fixation period, a sinusoidal grating, was presented either in the left or right hemifield and the participants had to discriminate its orientation (valid trial in this example). (D) *Example trial of the detection task*. At the beginning of each trial, a bullseye fixation point was shown. Then, a low-contrast sinusoidal grating was presented in the left, right or both hemifields. Participants were instructed to indicate the location of the stimulus and the contrast of the stimulus was adapted according to the participant’s performance using a staircase procedure.

In the framework of the synchronization theory [21,49], we assumed that tACS at IAF, but not at IAF+2Hz, IAF-2Hz, stimulates in phase with the intrinsic dominant alpha frequency and thereby progressively enhances intrinsic alpha power (Fig 2A). We therefore predicted that only left posterior parietal tACS at IAF, but not at IAF+/-2Hz, induces a leftward lateralization of alpha power relative to sham, which would in turn also result in a visuospatial attentional leftward bias (Fig 2B). Furthermore, we anticipated an association between the neural and the behavioral stimulation effect across participants. Accordingly, participants with a strong tACS-induced lateralization of alpha power were expected to display a stronger visuospatial attentional leftward bias and vice versa. Our results revealed an alpha power lateralization effect in the IAF but not in the IAF+/-2Hz condition, with a leftward lateralization of alpha power relative to sham. Only for the IAF stimulation condition, we also found that a high value of leftward alpha power lateralization co-occurred with a leftward visuospatial attention bias, which differed from the sham condition. Moreover, the magnitude of the alpha power lateralization effect predicted the visuospatial attention bias in the IAF stimulation condition.

**Figure 2.**
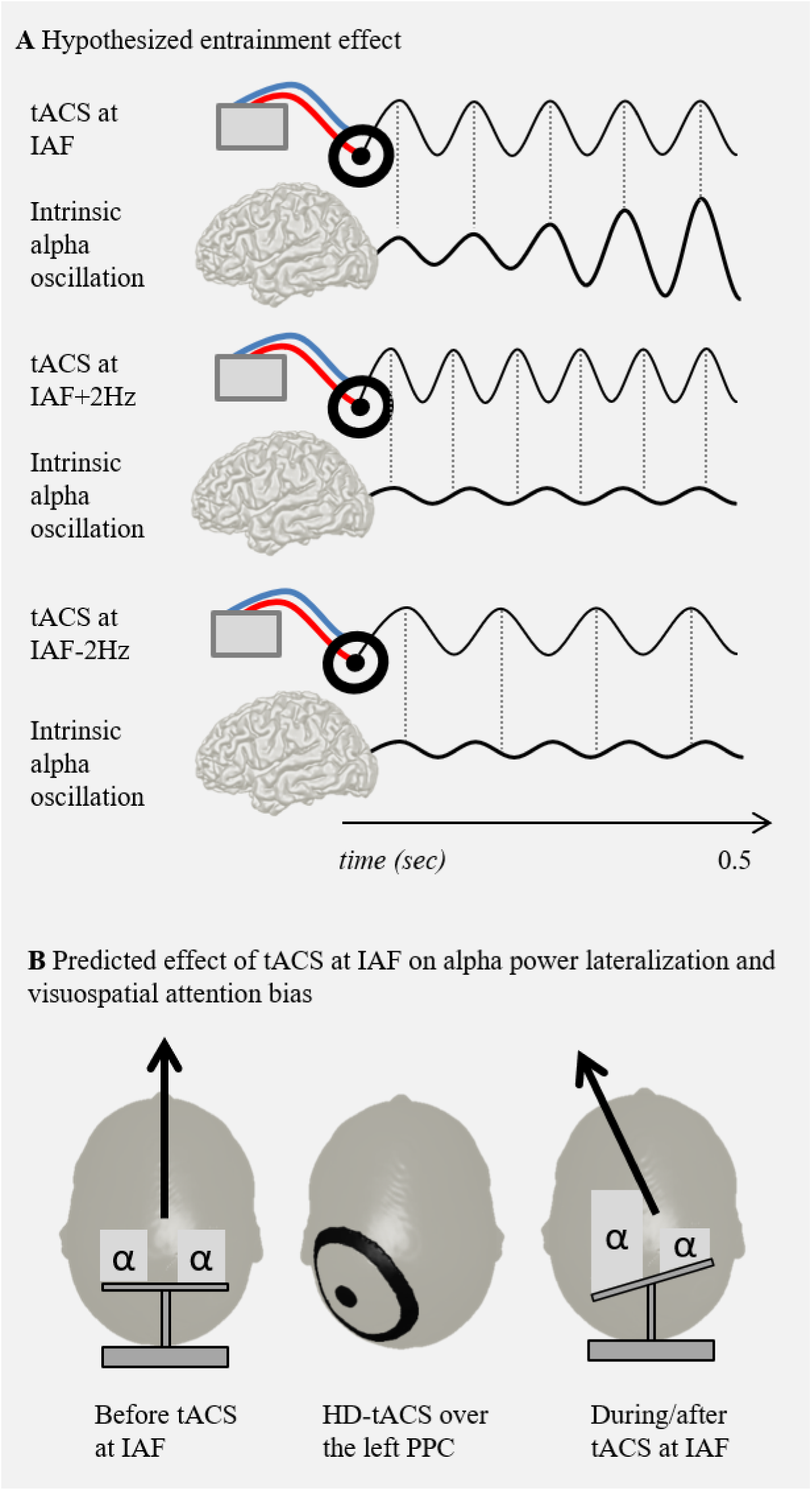
Rationale and hypotheses. (A) *Proposed effect of tACS at IAF, IAF+2Hz and IAF-2Hz on the intrinsic alpha power*. Depicted are the alternating currents of tACS at IAF, IAF+2Hz and IAF-2Hz respectively as well as the expected effect of the alternating current on intrinsic alpha power over time. The dashed lines indicate the phase of the tACS current relative to the phase of the intrinsic alpha oscillation. While tACS at IAF boosts intrinsic alpha power by consistently stimulating in phase, the currents of tACS at IAF+/-2Hz are not synchronized to the intrinsic alpha oscillation and therefore do not induce the same increase in alpha power. (B) *Predicted effect of tACS at IAF on alpha power lateralization and visuospatial attention bias*. We hypothesized that before tACS at IAF, alpha power is equal in both hemispheres, accompanied by an unbiased visuospatial attentional locus. After tACS at IAF over the left posterior parietal cortex (PPC), alpha power lateralizes to the left hemisphere, which results in a visuospatial attentional leftward bias.

## Results

### EEG data

#### IAF is a stable trait marker and negatively correlates with age

To verify the reliability of the IAF, we first examined its between-subject and within-subject variation. Our results show that the IAF spanned the 8 to 11.4Hz range across participants and negatively correlated with age (r_19_ = -.573, p = .007) (Fig S1). The test-retest reliability between the IAF estimates of the four sessions was very high as indicated by an average measure intraclass correlation coefficient (ICC) of .98 (F(20, 60) = 50.45, p < .001). This shows that the IAF is a stable trait marker, with minimal variation between sessions.

#### tACS at IAF but not at IAF+/-2Hz induces a leftward lateralization of alpha power

The effect of tACS on alpha power lateralization was quantified with the alpha power lateralization index (ALI), which indicates the percentage increase in alpha power (from the pre-to the post-measurement) for the left relative to the right hemisphere. Figure 3A depicts ALI per stimulation condition and the pattern of results suggests that tACS at IAF induced a leftward lateralization of alpha power. To test these data trends, we fitted a mixed model on the electrophysiological entrainment index ALI using stimulation condition as a factor. We found a significant main effect stimulation condition (F_3,60_ = 2.91, p = 0.042) and planned comparisons between the active stimulation conditions and sham revealed a difference between the IAF tACS and sham condition with a p-value equal to .05 (t_60_ = 2.29). There was a higher ALI for the IAF (M=8.16, SE=6.98) as compared to the sham condition (M=-11.46, SE = 6.98) indicating a leftward lateralization of alpha power in the IAF relative to the sham condition. The two other comparisons did not reach significance (IAF+2Hz vs sham: t_60_ = -.24, p = 1.00; IAF-2Hz vs sham: t_60_ = 1.46, p = .30) (Fig 3A). Hence, only tACS at IAF but not at IAF+/-2Hz induced a leftward lateralization of alpha power relative to sham. Note that this analysis focused on the first minute of the post-measurement to maximise entrainment effects. An analysis of the full three minutes of EEG data led to a similar pattern of results but no significant effects (Fig S2A).

**Fig 3.**
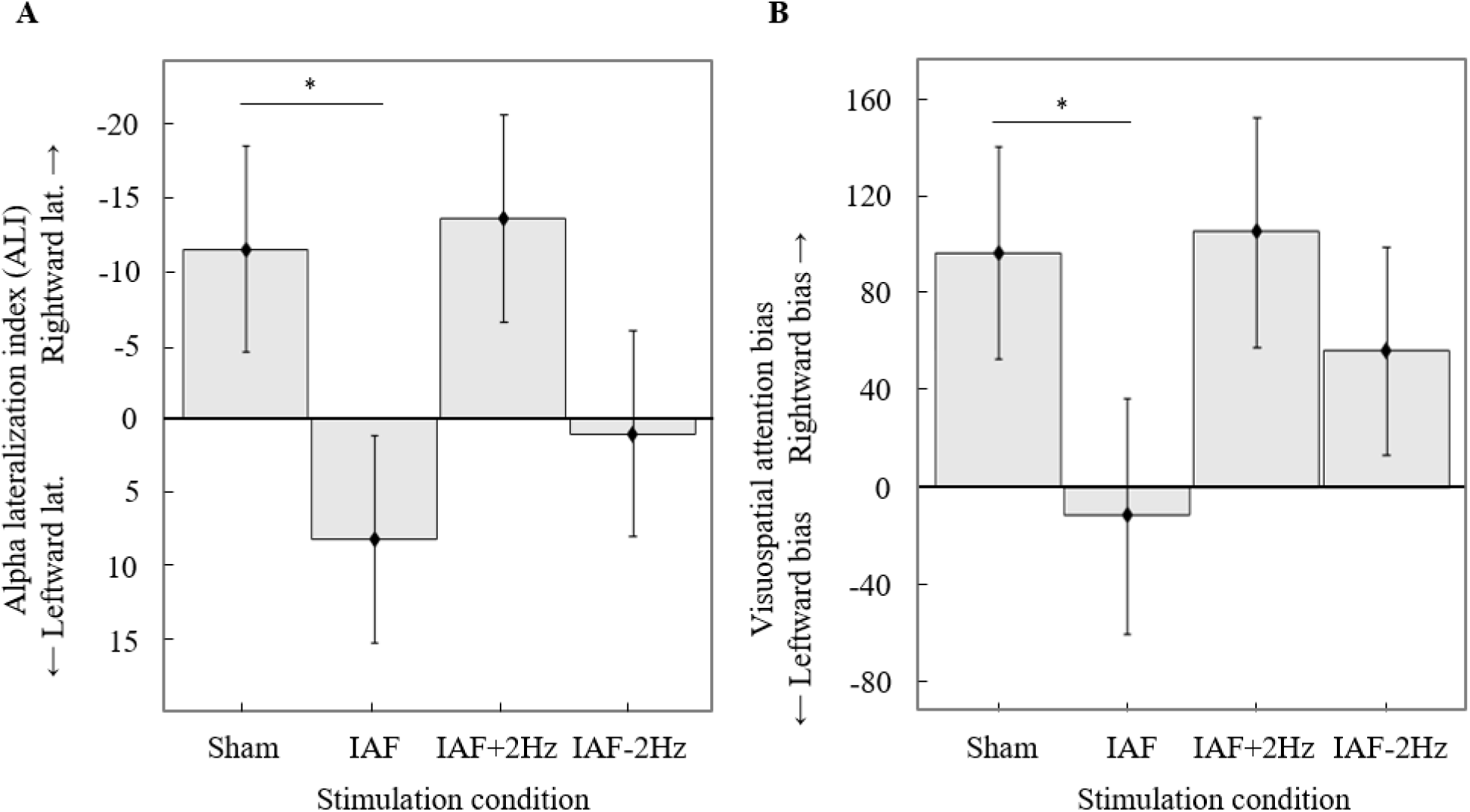
Electrophysiological and behavioral stimulation effect. (A) *Alpha lateralization index (ALI) per stimulation condition*. A positive value of ALI indicates a greater increase in alpha power in the stimulated left relative to the non-stimulated right hemisphere. (B) *Spatial cueing task: Visuospatial attention bias per stimulation condition for a high value of the covariate ALI*. A positive value of the visuospatial attention bias means that participants were less efficient in responding to target stimuli in the left relative to stimuli in the right hemifield. There was a significant interaction between stimulation condition and the covariate ALI. Here, the visuospatial attention bias per stimulation condition is depicted for a high value of the covariate (results of a simple slope analysis). The lines and asterisks indicate the results of planned comparisons between sham and the active stimulation conditions and mark p-values ≤ .05. The error bars depict the pooled standard error of the mean across participants.

To conclude, only tACS at IAF but not at IAF+/-2Hz induced a leftward lateralization of alpha power relative to sham. Further analyses per hemisphere yielded ambiguous information as to whether the ALI changes were due to power changes in the left or right hemisphere (Fig S3).

#### tACS at the control frequencies does not affect power at and around the stimulation frequency and does not shift the intrinsic IAF

We also assessed the effects of tACS on the lateralization of power at and around the stimulation frequency (PLI) to find out whether tACS at IAF-2Hz and IAF+2Hz modulated lower and upper alpha power respectively. For this, we directly compared tACS at IAF+/-2Hz to tACS at IAF and sham by fitting a mixed model on PLI with stimulation condition (all four conditions) as factor. We found a main effect of condition (F_3,60_ = 3.01, p = .037), driven by a significant higher PLI score in the IAF (M = 8.16, SE = 6.22) as compared to the sham condition (M = -11.46, SE = 6.22) (t_60_ = 2.44, p = .034). All other comparisons were not significant (IAF+2Hz vs sham: t_60_ = .03, p = 1.00; IAF-2Hz vs sham: t_60_ = 1.70, p = .19) (IAF+2Hz: M = -11.25, SE = 6.22; IAF-2Hz: 2.17, SE = 6.22) (Fig S4). This means that only tACS at IAF, but not at IAF+/-2Hz, induced a leftward lateralization in power at and around the stimulation frequency relative to sham.

In another control analysis, we tested whether tACS at the flanking control frequencies IAF+/-2Hz shifted the IAF towards the stimulation frequency. To this end, we analysed the change in IAF from the pre-to the post-measurement and fitted a mixed model on the IAF change score with stimulation condition as factor. There was no significant effect (F_3,60_ = .48, p = .699) (IAF: M = -.17, SE = .07; IAF+2Hz: M = -.11, SE = .07; IAF-2Hz: M = -.22, SE = .07; sham: M = -.19, SE = .07) indicating that tACS had no effect on the intrinsic IAF (Fig S5).

To conclude, we found an effect of tACS at IAF on alpha power lateralization but no effect of tACS at IAF+/-2Hz on alpha power, power at and around the stimulation frequency or intrinsic IAF. This underlines the frequency-specificity of the electrophysiological stimulation effect.

### Behavioral data

#### Spatial cues modulate task performance

The average accuracy in the spatial cueing task was 90% and ranged between 61%-100%. First, we assessed the cueing effect by looking into the differences in reaction time (RT) between the three different type of cue trials. We used the data of the sham condition for this analysis. We found a main effect of type of cue (F_2, 38.01_ = 9.47, p < .001). Accordingly, participants responded faster in valid (M = 593.67, SEM = 40.39) as compared to neutral (M = 612.83, SEM = 40.41) (t_38.02_ = -3.13, p = .003) and invalid trials (M = 619.31, SEM = 40.41) (t_38.02_ = -4.18, p < .001). The difference between invalid and neutral trials turned out to be not significant (t_38.00_ = 1.06, p = .297). This means that the valid but not the invalid cues modulated RTs. In the next sections, we investigated (for all three cue types) whether left tACS caused an attentional advantage for the processing of targets presented in the left hemifield. tACS at IAF, but not at IAF+/-2Hz induces a visuospatial attentional leftward bias in the spatial cueing task

To verify whether tACS at IAF induced a visuospatial attentional leftward bias, we fitted a mixed effect model on the visuospatial attention bias score (inverse efficiency_left targets_ – inverse efficiency_right targets_) using stimulation condition and type of cue as factors and ALI as covariate. There was a significant interaction effect between stimulation condition and ALI (F_3, 200.84_ = 3.01, p = .031). All other main and interaction effects were not significant (stimulation condition: F_3, 199.10_ = 1.24, p = .298; type of cue: F_2, 198.05_ = 1.23, p = .294; ALI: F_1, 208.70_ = .88, p = .349; stimulation condition × type of cue: F_6, 198.05_ = .45, p = .847; type of cue × ALI: F_2, 198.07_ = .35, p = .705; stimulation condition × type of cue × ALI: F_6, 198.10_ = .31, p = .931). This means that the effect of stimulation condition on visuospatial attention bias depends on the electrophysiological stimulation effect ALI. A simple slope analysis revealed that the stimulation condition effect was significant for a high (F_3, 200.10_ = 3.90, p = .002) but not for a low (F_3, 200.05_ = 2.45, p = .130) or average value (F_3, 199.26_ = 1.38, p = .500) of the covariate ALI. Accordingly, there is an effect of stimulation condition on the visuospatial attention bias when the model assumes a high value but not a low or average value of the covariate ALI. For the high value of the covariate ALI, the IAF stimulation condition (M = -11.83, SE = 48.77) differed from sham (M = 96.42, SE = 43.77) (t_200.51_ = -3.05, p = .006), in line with an increased visuospatial attentional leftward bias. All other comparisons were not significant (IAF+2Hz vs sham: t_199.25_ = .27, p = 1.00; IAF-2Hz vs sham: t_199.95_ = -1.45, p = .296) (Fig 3B; see also Fig S6 for the effect per cue type and Fig S2B for the behavioral effect including ALI with all three minutes of the EEG post-measurement as a covariate). This indicates that only tACS at IAF but not at IAF+/-2Hz induced a significant leftward shift of visuospatial attention relative to the sham condition for a high value of the covariate ALI. Further analyses per target location could not reveal whether the visuospatial attention bias effect was caused by performance changes in left or right target location trials (Fig S7).

#### In the IAF stimulation condition, the electrophysiological stimulation effect predicts the behavioral stimulation effect

We also analyzed the association between the electrophysiological and the behavioural stimulation effect in the spatial cueing task to further explore the significant interaction between stimulation condition and the covariate ALI on the visuospatial attention bias score. To this end, we ran linear regression analyses between ALI and the visuospatial attention bias score per stimulation condition. In the IAF stimulation condition, the electrophysiological stimulation effect ALI predicts the behavioural stimulation effect (*b* = -.57, p = .014) and, according to R^2^, explains 32% of the variance. For all the other stimulation conditions, there was no association between ALI and the visuospatial attention bias (IAF+2Hz: b = .19, p = .458; IAF-2Hz: b = -.20, p = .434; sham: b = .25, p = .289) (Fig 4). While the regression coefficient of the IAF stimulation condition differed from sham (t(199.56)= -2.48, p = .028), no differences between the control stimulation conditions and sham could be found (IAF+2Hz vs sham: t_199.03_ = .69, p = .984; IAF+2Hz vs sham: t_199.09_ = .04, p = 1.00). Note that also here, only the first minute of the post EEG measurement was included in the analysis. However, an analysis of the full three minutes EEG data lead to the same pattern of results as well as similar statistics (Fig S8).

**Fig 4.**
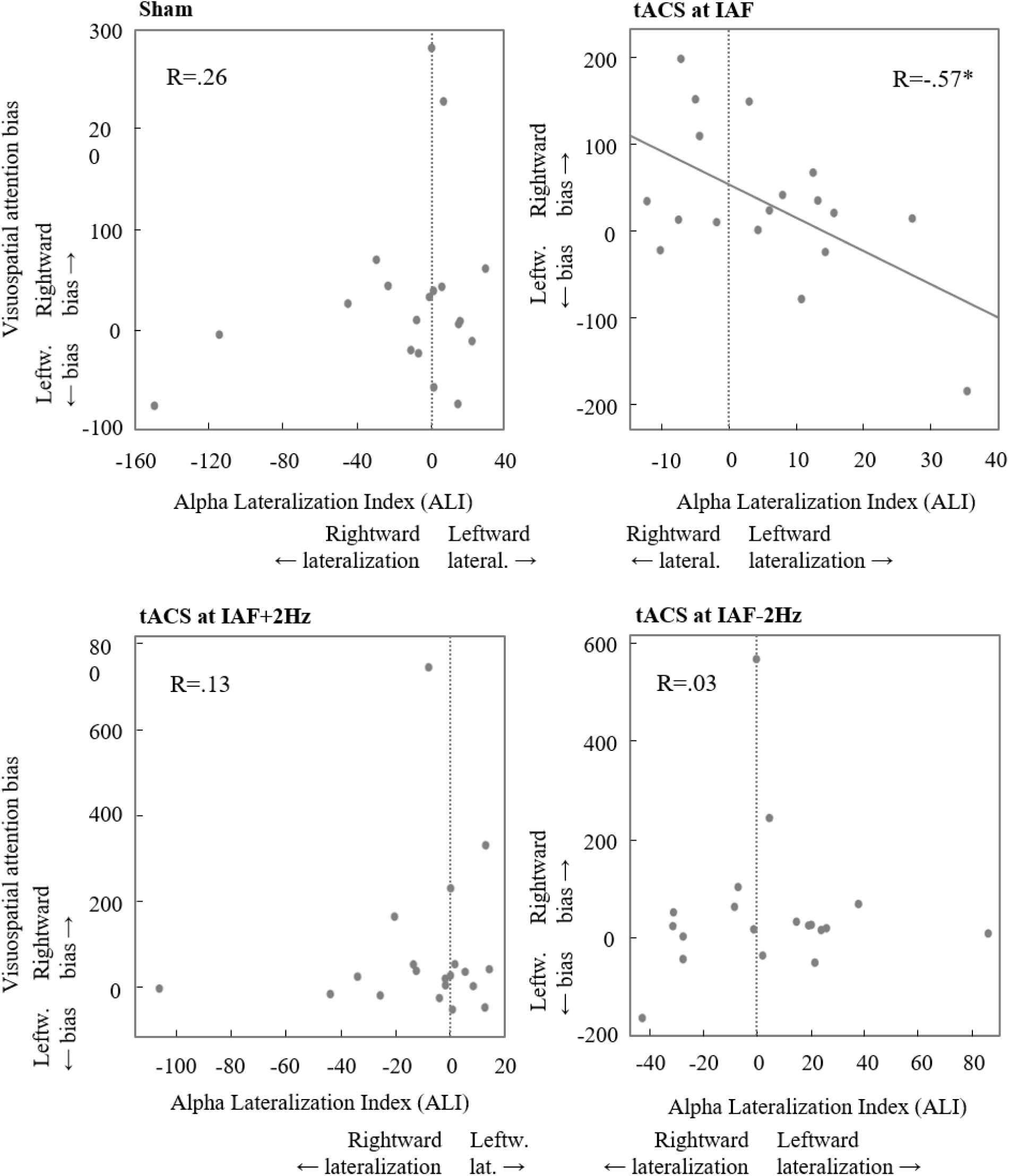
Association between ALI and visuospatial attention bias in the spatial cueing task per stimulation condition. A positive value of ALI indicates a greater increase in alpha power in the stimulated left relative to the non-stimulated right hemisphere from the pre-to the post-measurement. A positive value of the visuospatial attention bias score means that the participants were less efficient in responding to target stimuli in the left relative to stimuli in the right hemisphere. Per subplot, each point depicts the data of one participant. The asterisk marks effects with a p-values ≤ .05. The R-value indicates the regression coefficient.

To conclude, only in the IAF but not in the IAF+/-2Hz condition did the electrophysiological and the behavioural effect correlate in magnitude, with greater tACS-induced leftward lateralization of alpha power predicting greater visuospatial attentional bias to the left.

#### The stimulation effect does not depend on age or IAF

Subsequently, we tested whether age or IAF affected the behavioural stimulation effect by running linear regression models on the visuospatial attention bias score in the IAF stimulation condition. We found no effect of age (b = .30, p = .231) or IAF (b = .24, p = .330) on the visuospatial attention bias, which means that the behavioural stimulation effect is independent of those two factors. Furthermore, we tested whether age or the IAF had an influence on the association between ALI and visuospatial attention bias in the IAF stimulation condition. According to log likelihood tests, adding age and IAF as predictors did not significantly improve the model (model_ALI_ vs model_ALI, IAF_: X2(1, N = 18) = 1.70, p = .19; model_ALI_ vs model_ALI, age_: X2(1, N = 18) = .00, p = 1.00; model_ALI_ vs model_ALI, IAF, age_: X2(2, N = 18) = 4.17, p = .12). This means that a model with only ALI as predictor explains the data best.

#### tACS does not affect visual detection performance

To find out whether the tACS effects on visuospatial attention performance are task specific, we performed a mixed model analysis on the contrast threshold bias score of the detection task using stimulation condition as factor and ALI as covariate. There were no significant main (stimulation condition: F_3, 51.41_ = .41, p = .746; ALI: F_1,60.81_ = 1.11, p = .297) or interaction effects (F_3, 54.73_ = .14, p = .934) (Fig 5). Additionally, we ran the same mixed model analysis using the bias in indicated target location score as dependent variable. Also here, the main (stimulation condition: F_3,50.05_ = 1.77, p = .166; ALI: F_1,55.19_ = .27, p = .607) and interaction effects (F_3, 51.71_ = .13, p = .944) turned out to be not significant (Fig S9). Consequently, tACS did not significantly affect detection performance in our experiment, which shows that the effect of tACS at IAF is task specific.

**Fig 5.**
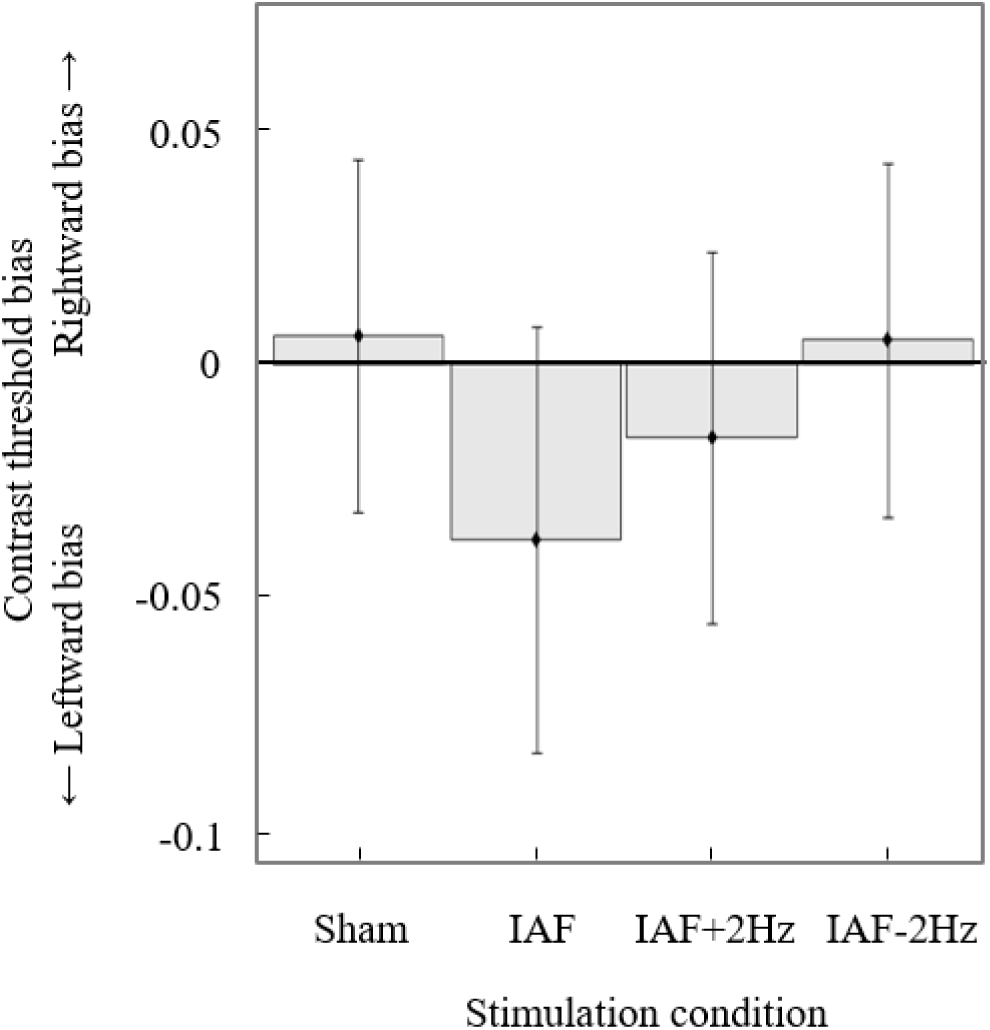
Detection task: Contrast threshold bias per stimulation condition. A positive value means that participants were able to detect lower contrast stimuli in the right relative to the left hemifield (rightward bias). The error bars depict the pooled standard error of the mean across participants.

#### A post-questionnaire confirms that blinding was effective

Information about the experimental hypotheses and stimulation conditions were withheld from the participants until completion of the experiment. At the end of each session we administered a questionnaire, which prompted the participants to evaluate whether real or sham stimulation was applied. According to the Wald chi square, the actual stimulation condition did not affect the rated stimulation condition (X^2^(3, N = 80) = 2.06, p = .56), which means that participants could not differentiate between the active and sham stimulation conditions and blinding was therefore effective.

## Discussion

In this experiment, we measured the effects of left posterior parietal HD-tACS at an individualized alpha stimulation frequency on occipitoparietal alpha power and visuospatial attention performance. We hypothesized that tACS at IAF but not at flanking frequencies induces a leftward lateralization of alpha power and a leftward bias in visuospatial attention. We also expected that the size of the behavioral stimulation effect would depend on the size of the neural stimulation effect, i.e. the greater the leftward lateralization of alpha power, the greater the leftward bias in visuospatial attention. In separate sessions and in a within-subject design, we applied HD-tACS at IAF, two control frequencies IAF+2Hz and IAF-2Hz and sham to the left PPC. During stimulation, we measured the visuospatial attention bias with a spatial cueing and a detection task and immediately before and after stimulation, we acquired EEG data to assess the changes in alpha power. To have a great variation in IAF and to be able to generalize our findings to a wider target group, we tested participants of various age groups spanning from adolescence to mature adulthood. Our data show a leftward lateralization of alpha power in the IAF stimulation condition, which differed from sham. Moreover, for a high value of alpha power leftward lateralization, there was a visuospatial attentional leftward bias in the IAF stimulation condition, which significantly differed from sham. Only for the IAF stimulation condition, we also found the hypothesized link between the electrophysiological and the behavioural stimulation effect across participants: the greater the leftward lateralization of alpha power, the greater the leftward bias in visuospatial attention in the spatial cueing task. Hence, there is a behavioral stimulation effect for those participants who also display a strong neural stimulation effect. No alpha power lateralization effect or visuospatial attention bias effect were induced by tACS at the control frequencies (IAF+/-2Hz). Further analyses revealed that the behavioural stimulation effect and the link between the electrophysiological and the behavioural stimulation effect were independent of age and intrinsic IAF. To conclude, our results show a frequency-specific effect of tACS at IAF on visuospatial attention performance and oscillatory alpha power as well as a link between the neural and the behavioural stimulation effect. This suggests that alpha oscillations play a causal functional role in the modulation of visuospatial attention and shows that tACS can be used to modulate it. The frequency specificity of the effects supports the synchronization theory as the neural basis of the behavioural and oscillatory stimulation effects and suggests that an individualization of the stimulation frequency is necessary in heterogenous target groups with a wide variation in IAF.

### Frequency specific stimulation effect and personalized stimulation protocols

In addition to tACS at IAF, we included two control frequency stimulation conditions (tACS at IAF+/-2Hz) in our experimental design to test the frequency specificity of the stimulation effect. While it has been proposed that tACS operates via synchronization of the intrinsic oscillation to the applied alternating current [21], some of the underlying assumptions of the synchronization theory have not been verified yet for tACS. The synchronization theory proposes that the stimulation effect diminishes with increasing deviation of the stimulation frequency from the intrinsic frequency [21,49]. However, we are not aware of any experiment that compared the effect of tACS at the intrinsic frequency to tACS at close flanking control frequencies. Here we found a frequency-specific effect of tACS at IAF on alpha power lateralization and visuospatial attention bias as well as a direct link between those two dependent variables. In contrast, tACS at the flanking control frequencies IAF+/-2Hz did not modulate alpha power lateralization or the visuospatial attention bias, in line with the synchronization theory.

The Arnold tongue depicts the degree of synchronization as a function of two factors: the amplitude of the driving force and Δfrequency, which is the difference in frequency between the driving force and the intrinsic oscillator [49]. The weaker degree of synchronization at larger Δfrequency can be overcome by increasing the amplitude of the driving force, which increases the Δfrequency window in which a synchronization effect can be observed. Notbohm and colleagues [50] verified these assumptions for rhythmic sensory entrainment by examining the effect of light flashes at various frequencies and intensities on phase synchronization. Only in the rhythmic but not in an arrhythmic stimulation condition did they find greater phase coupling with increasing stimulation intensity and smaller Δfrequency. Future research should systematically assess the effect of various tACS stimulation intensities and levels of detuning between IAF and stimulation frequencies on EEG to verify whether a similar pattern exists for tACS.

The frequency-specificity of the stimulation effects in the present experiment is not only theoretically relevant but has also important practical implications for the experimental and clinical use of tACS. The effect of tACS on alpha power lateralization and visuospatial attention bias was limited to stimulation at IAF, which suggests that an individualization of the stimulation frequency might be necessary in heterogenous samples with a wide variation in IAF. As the IAF negatively correlates with age in adults [51,52] and is particularly low in patients with dementia [53–55], traumatic brain injury [56,57] or stroke [58,59], an individualization of the stimulation frequency might be especially important for the clinical use and the application of alpha tACS in a sample covering a wide age range. Furthermore, tuning tACS to the IAF instead of stimulating at a fixed frequency of 10Hz, might reduce variability [42,60,61] and therefore lead to more robust tACS effects even in homogenous samples of young, healthy participants.

### Link between electrophysiological and behavioral stimulation effect

Our data reveal a link between the behavioral and electrophysiological stimulation effect in the IAF stimulation condition: participants with a high alpha power lateralization effect also displayed a marked visuospatial attentional leftward bias and vice versa. In an EEG paradigm, Thut and colleagues [15] found a similar association between alpha power lateralization over occipitoparietal sites and the visuospatial attention bias in an attentional cueing task. Collapsed over trials, there was a leftward lateralization of alpha power, in accordance with an overall attentional leftward bias and on a trial-by-trial level, alpha power lateralization predicted the reaction time to upcoming liminal stimuli in the left or right hemifield. Here, we significantly extend those findings by demonstrating a correlation in magnitude between the tACS-induced changes in alpha power lateralization and the tACS-induced visuospatial attention bias. Furthermore, the alpha power lateralization effect in the IAF tACS condition predicted the visuospatial attention bias and explained 32% of its variance. This suggest that alpha power plays a causal functional role in the modulation of visuospatial attention and shows that the magnitude of the electrophysiological tACS effect is a determinant for the magnitude of the behavioral tACS effect.

The association between the magnitude of the electrophysiological and the behavioral stimulation effect also had practical implications for the statistical analysis of the behavioral data. The electrophysiological stimulation effect was included as a covariate in a mixed model analysis to explain variance and to analyze the behavioral effect at different levels of the electrophysiological stimulation effect. In contrast to an analysis based on a dichotomous categorization into responders and non-responders, this mixed model approach does not require a division into small subgroups. Instead, the regression equation takes the data of all participants into account and predicts the stimulation effect at different values of the covariate based on the regression equation (simple slope analysis). This form of analysis could be useful for various brain stimulation studies suffering from variable, weak or inconsistent stimulation effects [62,63,72–75,64–71] or studies that want to elaborate on the association between the electrophysiological and the behavioral stimulation effect. Another advantage of mixed model analysis is that the omission of observations due to outlier removal, technical faults or data loss does not lead to exclusion of data on a subject level. Instead, only single cells are omitted, while the rest of the data is still included in the calculation of the regression equation.

### Task specific tACS effect

Similarly to previous tACS and tDCS studies that targeted the left posterior parietal cortex, we found a behavioral stimulation effect in the spatial cueing but not in the detection task [43,76]. In the spatial cueing task, participants have to discriminate the orientation of left and right target stimuli after symbolic, central cues instructed them to direct attention towards either hemifield. This task therefore assesses higher-level attentional processes such as the ability to perform endogenous (top-down) attention shifts as well as the efficiency in discriminating the orientation of lateralized target stimuli. In contrast, the detection task measures the ability to detect liminal stimuli in the left, right or both hemifields by determining contrast thresholds through staircase procedures. It therefore assesses low-level visual processing and the ability to perform exogenous attention shifts towards lateralized stimuli.

There are various explanations that could account for the task specific stimulation effect in this experiment. EEG studies that employed spatial cueing tasks found a lateralization of alpha power after presentation of a central, symbolic endogenous cue, but before a lateralized target stimulus was shown [12–15]. This suggests that alpha power changes are associated with endogenous attention shifts in a spatial cueing task. Furthermore, we applied a HD-tACS ring electrode montage over the posterior parietal cortex, an area that includes a key node of the dorsal attention network (DAN) - the intraparietal sulcus (IPS). The DAN is involved in endogenous shifts of attention, which is in line with the effect of posterior parietal tCS on spatial attention performance in endogenous cueing tasks. Moreover, it has been proposed that voluntary orienting is dissociated from target detection in the posterior parietal cortex. An fMRI study showed that the posterior parietal cortex is involved in voluntary attention shifts while target detection is rather regulated by the temporoparietal junction [77]. All of this might account for the stimulation effect in the spatial cueing task and the absence of effects in the detection task (however, see [78,79]).

### High-density electrode montage and stimulation site

The effect of unilateral HD-tACS at alpha frequency on visuospatial attention is in line with recent alpha tACS research on visuospatial [43,44] and auditory spatial attention [45,46]. However, previous tACS experiments found no or inconsistent effects of lateralized alpha tACS on the visuospatial attention bias [65,75]. These inconsistent results are surprising considering the quite established association between alpha power lateralization and visuospatial attention bias [12–15,18].

One aspect that distinguishes the experiments with an effect of tACS on the visuospatial attention bias [43,45,46] from experiments that report no or inconsistent effects [65,75], is the high-density ring electrode montage (however, see [44]). In contrast to traditional tACS montages, consisting of two rectangular or round tACS electrodes, the ring electrode montage creates a more focused electrical field between the large ring and the small disk electrode [80]. This restricted current flow allows for a more precise anatomical targeting limited to one hemisphere, which is confirmed by a current simulation for the electrode montage in this experiment (Fig 1B), which shows that the electrical field was confined to the left posterior parietal cortex.

An alternative explanation for the previous inconsistent and null results lies in the stimulation site. While left hemispheric stimulation tACS at alpha frequency modulated the visuospatial attention bias [43,44], no such effect could be found for right hemispheric stimulation [44,65,75]. In line with this, various EEG studies on auditory and visuospatial attention reported stronger alpha power dynamics over the left hemisphere [14,81–83]. Furthermore, Meyer and colleagues [84] showed that the task-related fMRI signal during a spatial cueing task is greater in the left as compared to the right frontoparietal attention network, especially after presentation of an invalid spatial cue. The left hemisphere also displayed a greater change in functional connectivity during a spatial cueing task as compared to the right hemisphere. In the right hemisphere, the functional connectivity was tonically higher during rest, while the left hemisphere was more specifically recruited during a condition of high attentional demands. This might explain why left but not right parietal tACS modulated visuospatial attention performance.

### Clinical relevance, duration of after-effects and ideas for future research

Our findings suggest a potential clinical application in which unilateral tACS at IAF is applied to correct the pathological attentional rightward bias of hemineglect patients. Clinical trials will show whether this stimulation protocol leads to similar leftward shifts of visuospatial attention in heminglect patients. As tACS is a noiseless, easy-to-use and portable technique, it could be applied simultaneously with standard therapies such as visual-spatial scanning and figure description training [85–87] in the ambulatory care to enhance treatment effects. Furthermore, tACS has been proposed to induce neuroplastic changes under the stimulation site [19,42,88], which means that tACS could represent a promising treatment option with long-term benefits.

However, in this experiment we showed that a single application of left posterior parietal tACS for 40 minutes at a low intensity of 1.5mA might not suffice for inducing long-lasting effects. Previous research demonstrated that alpha tACS to the medial occipitoparietal cortex can enhance alpha power for up to 70 minutes after stimulation [89]. However, we are not aware of any prior study that verified whether a lateralized tACS montage leads to a similar modulation of alpha power. Here we showed for the first time (to our knowledge) that left posterior parietal tACS at IAF induces a lateralization of alpha power towards the stimulation site. In contrast to previous research with medial montages, however, this effect did not last for up to 70 minutes [89], but could only be shown for the first minute of the post-measurement. An analysis of the full three-minute post-measurement data revealed no significant differences in alpha power lateralization between the IAF and sham stimulation condition, which suggests that the after-effects of the tACS intervention weaken or fade away. Hence, the oscillatory effect of a lateralized tACS montage is not necessarily comparable to central montages, which highlights the necessity of combining tACS with EEG in case of previously untested montages in order to verify that tACS leads to the expected changes in oscillatory power. However, it is important to note that the correlation between the alpha power lateralization effect and the visuospatial attention bias in the IAF stimulation condition was significant for all three minutes, which strengthens the fundamental effects revealed for the first minute. To enhance the clinical applicability of the tACS interventions with a laterally placed setup, further research will be necessary to find strategies for prolonging the after-effects.

It is plausible that during the long period of rest in the post-measurement, participants adjusted the tACS-induced visuospatial attentional leftward bias and the associated alpha power lateralization by directing the spatial attention locus back to the midpoint. Alternatively, interhemispheric dynamical interactions might have pulled the alpha power lateralization back into its original balanced state. The interhemispheric inhibition theory claims that the two hemispheres mutually inhibit each other via transcallosal connections [90–93]. These transcallosal connections might have regulated the adjustment of the interhemispheric alpha power imbalance. Additionally or alternatively, a single application of left posterior parietal tACS for 40 minutes at a low intensity of 1.5mA might not suffice for inducing long-lasting neuroplastic changes. Mohsen and colleagues [94] showed that a repeated application of transcranial random noise stimulation (TRNS) in chronic tinnitus patients enhances stimulation effects. Similarly, a repeated application of left posterior parietal tACS at IAF could amplify and prolong the after-effects by inducing accumulative neuroplastic changes. Furthermore, Nitsche and Paulus [95] showed that the duration of the after effects on motor cortex excitability following transcranial direct current stimulation (tDCS) increase with stimulation intensity and stimulation duration. In the same way, Vöröslakos and colleagues [96] demonstrated that the effect of intersectional short pulse stimulation on alpha power depends on stimulation intensity. Hence, future research should investigate whether this can be translated to tACS, e.g. whether higher stimulation intensities, a repeated application of the stimulation protocol and longer stimulation durations lead to stronger and longer-lasting (after-) effects.

## Conclusion

We hereby demonstrate a causal and frequency-specific effect of tACS at IAF on visuospatial attention bias and alpha power lateralization. The visuospatial attention bias effect was underpinned by and directly linked to the alpha power lateralization effect. This suggests that alpha oscillations play a causal functional role in the modulation of visuospatial attention and demonstrates that tACS can be used to modulate it. We also present a statistical approach, which allows for an analysis of the behavioral effect at different levels of the electrophysiological effect. This approach could be useful for future brains stimulation studies that suffer from weak or variable stimulation effects or simply want to elaborate on the association between the electrophysiological and the behavioral stimulation effect. The frequency specificity of our findings, limited to tACS at IAF, is in line with the theory of synchronization and suggests that an individualization of the stimulation frequency is necessary in heterogenous groups of participants with a wide variation in IAF. This might account for previous inconsistent or variable tACS effects and could be relevant for potential clinical applications of tACS in hemineglect patients.

## Methods

### Participants

We tested 21 healthy, right-handed volunteers (8 female, mean age (SD) = 45.38 (17.10) years, age range = 19-72 years) with normal or correct to normal vision. Participants filled in an informed consent and a tACS safety screening form prior to each session. In the safety screening form we scanned for e.g. neurological disorders, skin diseases, medication and pregnancy, taking the recommended procedures of Antal and colleagues [97] into account. This study was performed in accordance to the Declaration of Helsinki and was approved by the Ethics Review Committee Psychology and Neuroscience (ERCPN) of Maastricht University (ERCPN number: 129). Participants received vouchers as compensation for their participation.

### Procedure

Each participant received active tACS at IAF, IAF+2Hz and IAF-2Hz as well as sham tACS on separate days and randomized order and the same procedure was followed in every session. To avoid carry-over effects, we scheduled sessions at least two days apart. Initially, participants performed a shortened practice version of the spatial cueing and the detection task. Then, we mounted recording EEG electrodes as well as stimulating tACS electrodes on the participant’s head. Before stimulation, three minutes of resting state EEG data were collected while participants kept their eye closed. This pre-measurement served as an estimation of alpha power before stimulation and was also used to determine the IAF. In order to account for potential day-to-day variations in the intrinsic frequency, the tACS stimulation frequency was always based on the IAF that was determined in the same session. Subsequently, tACS was applied at either IAF, IAF+2Hz, IAF-2Hz or sham while participants performed a spatial cueing and a detection task in randomized order. After completion of the tasks or after a maximum stimulation duration of 40 minutes, the tACS stimulator was switched off and three minutes of resting state EEG data were measured again (post-measurement) (Fig 1A). During the spatial cueing task, we recorded eye movements with an eye tracker. These data were used for offline analysis of the behavioural performance. Information about the experimental hypotheses and the applied stimulation protocol were withheld from the participants until completion of the experiment. To verify whether blinding was maintained, we administered a questionnaire at the end of each session, which prompted the participants to evaluate the stimulation condition based on the subjective experience.

### Eye tracker

An eye tracker (Eyelink1000, SR Research, Mississauga, Ontario, Canada) was used during the spatial cueing task. At the beginning of each session, we performed a 9-point calibration and validation procedure. Then, we assessed the participants’ gaze position sample by sample point using monocular eye tracking at 1000Hz. These data were used for the offline analysis. We did not record eye tracking data during the detection task because this task did not include an orienting or reorienting component.

### tACS and electric field simulation

We mounted a high-density ring electrode tACS montage (NeuroConn, Ilmenau, Germany) over the left posterior parietal cortex with a small circular electrode (Diameter: 2.1cm; Thickness: 2mm) positioned over P3 and a large ring electrode (Outer diameter: 11cm; Inner diameter: 9cm; Thickness: 2mm) centered on it. The international 10-20 EEG system was used to determine the electrode position P3 on the participants’ head. tACS was applied via an DC-stimulator plus (NeuroConn, Ilmenau, Germany) at a stimulation intensity of 1.5mA peak to peak. In the active tACS conditions, the stimulation frequency was tuned to IAF, IAF+2Hz or IAF-2Hz and the ramp up was set to 100 cycles. The tACS device was switched off as soon as the participant finished the task but never exceeded 40 minutes. Overall, the stimulation duration varied between 35-40 minutes. In the sham condition, we also applied tACS at IAF but ramped up and then immediately ramped down the stimulation with each 100 cycles. That way, we imitated the initial skin sensations of real tACS while minimizing neural and behavioral stimulation effects. Conductive gel (Ten20 paste, Weaver and Company, Aurora, CO, USA) was used to fasten the tACS electrodes on the skin and to keep impedances below 10kΩ.

For this experiment, we used a high-density ring electrode tACS montage to enable spatially focal stimulation [80] of the left PPC. An electric current simulation was performed to visualize the stimulated regions using a custom-written MATLAB script [36] interfacing with the software SimNIBS [98,99] (Fig 1B). For this simulation, we used a freely available individual head model of a healthy brain as an example participant [100] and modelled the electrodes with a random connector location. The conductivity of the ten20 paste was set to 8 S/m, an estimation based on the concentration of CI-in the gel [101].

### EEG apparatus and data acquisition

First, we marked the electrode positions P5, PO3, P6 and PO4 on the participants’ head according to the international 10-20 system. Then, single EEG electrodes were mounted at the marked spots using ten20 conductance paste (Weaver and Company). Spectral EEG was recorded via a BrainAmp MR Plus EEG amplifier (BrainProducts GmbH, Munich, Germany) and Ag-AgCl electrodes (BrainProducts GmBh, Munich, Germany). The recordings were online referenced to the left and offline re-referenced to both mastoids and the ground electrode was positioned over the right forehead. Impedances for all electrodes were kept below 5 kΩ and a sampling rate of 500Hz and a bandpass filter of 0.1-200Hz was used for online recording.

### Task description

The spatial cueing task was a classical Posner task including endogenous cues and was used to assess the participants’ speed and accuracy in discriminating the orientation of lateralized target stimuli in the left or right hemifield. Throughout the task, participants had to fixate on a central white fixation point, surrounded by a black or grey donut-shaped area, which was delimited by a black circle. A trial started with a jittered interval of 800-1200ms, during which only the white fixation point, surrounded by a grey area was presented. Subsequently, the grey area turned black for 500ms. Then, a central symbolic cue, which consisted of arrowheads pointing to the left (<<·<<), right (>>·>>) or both sides (<<·>>) flanking the central fixation point, was shown for 100ms. The directional cues (left or right arrow heads) predicted the correct target location with 80% validity. The cue was followed by a cue-target interval of 500ms during which only the central fixation point was presented. Then, the target stimulus was shown for 100ms in either the left or right hemifield at 7° eccentricity from the fixation point. This target stimulus consisted of a sinusoidal grating with a Gaussian envelope (spatial frequency = 1.5 cycles per degree, envelope standard deviation = 0.75°) and was rotated clockwise or counter clockwise by 45°. Participants were instructed to differentiate the orientation of the stimulus as fast and accurately as possible, pressing the numerical button 1 or 2 for counter clockwise and clockwise rotated stimuli respectively (Fig 1C). Trials with a very slow (>1000ms) or anticipatory (<120ms) response were repeated. The endogenous task took approximately 20 minutes and comprised 335 trials, of which 192 were valid, 48 invalid and 96 neutral cue trials.

The detection task measured the participants’ ability to detect low-contrast target stimuli in the left, right or both hemifields. First, participants manually downregulated the contrast of bilaterally presented stimuli until they were barely visible. This contrast served as an initial value for the subsequent staircase procedure. A trial started with the presentation of a white fixation point, surrounded by a grey donut-shaped area, which was delimited by a black circle. After 1s, the grey area turned black for 500ms. Then, the target stimulus, a randomly oriented sinusoidal grating (spatial frequency = 1.5 cycles per degree, envelope standard deviation = 0.75°), was presented for 100ms in the left, right or both hemifields at 14° eccentricity from the fixation point. Participants had to indicate the stimulus location, pressing the numerical button 1, 2 or 3 for left, bilateral and right target location respectively. In case no stimulus was perceived, the participants had to withhold the response (Fig 1D). On a trial-by-trial basis, the contrast of the left, right and bilateral stimuli were independently adjusted according to the QUEST staircase algorithm [102] as implemented in the Psychophysics Toolbox [103] for MATLAB (prior standard deviation = 1, beta = 3.5, gamma = 0.01, delta = 0.01, aim performance = 50% detection rate). QUEST is a psychometric procedure which uses Bayesian statistics to predict the participant’s contrast threshold based on the detection performance in the preceding trials. We used the function QuestQuantile to compute the trial-by-trial stimulus contrast based on the maximum likelihood estimate of the threshold. QuestMean was used to calculate the final detection threshold. As the contrast threshold of left, right and bilateral stimuli were independently determined, the detection task consisted of three interleaved staircase procedures of each 40 trials resulting in a total amount of 120 trials. The detection task took approximately 10 minutes.

Both tasks were presented on a gamma-corrected liyama ProLite monitor at 60Hz. The background luminance and the video mode were set to 100cd/m^2^ and 1920×1080 respectively. Participants had to place their chin into a chin rest to assure a viewing distance of 57cm as well as a central and stable position of the head. We used the software application Presentation (NeuroBehavioural Systems, Albany, CA) for the presentation of the stimuli and recording of the behavioural response. The behavioural responses were recorded via a standard USB-computer keyboard and the participant always pressed the response button with the right hand.

### Preprocessing

#### EEG

Resting state EEG measurements were analysed offline using the FieldTrip toolbox [104] as implemented in MATLAB (MathWorks). We segmented the EEG data into 5-second epochs, resulting in a frequency resolution of 0.2Hz. Trials with an amplitude over time variance deviating more than 2 standard deviation from the mean were rejected and excluded from the subsequent analyses. Then, we ran a Fourier analysis using Hanning tapers to calculate the power spectra between 1 and 100 Hz per channel and participant. The IAF was computed by averaging the power values over time and all four occipitoparietal channels and identifying the peak frequency in the power spectrum between 7 and 13Hz.

The alpha lateralization index (ALI) served as a measure of the neural stimulation effect and was determined by subtracting the proportion increase in alpha power (PIA) in the right hemisphere from PIA in the left hemisphere. PIA was defined as follows

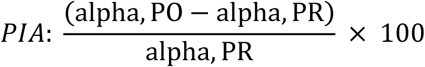

in which alpha reflects the average of the individual alpha power in the frequency interval IAF-1Hz to IAF+1Hz, for the pre (PR) and the post-measurement (PO) respectively. For the post-measurement, we only analysed the first minute of the post-measurement to maximize entrainment effects. The percentage increase in power (PIP) and the power lateralization index (PLI) were calculated in the same way as PIA and ALI respectively with the only difference that the frequency window for the analysis was not centred on the individual alpha frequency band but on the stimulation frequency. For the IAF and sham stimulation conditions, this means that we analysed the alpha power in the frequency window spanning from IAF-1Hz to IAF+1Hz. For the IAF+2Hz and IAF -2Hz condition, we derived the power for a lower (IAF-3Hz to IAF-1Hz) and higher (IAF+1Hz to IAF+3Hz) frequency band respectively.

#### Spatial cueing task

For the spatial cueing task, we removed trials containing eye blinks and eye movements within a window of 100ms before cue onset until stimulus onset exceeding 2° of visual angle (7% of all trials). For the analysis of the reaction time scores (RT) we excluded trials with an incorrect or missing response (10% of all trials) and with deviating RT scores, falling outside the median +/-1.5*interquartile range (IQR) per stimulation condition and trial type (2% of all trials). Subsequently, we calculated accuracy and RT scores per condition. As the distribution of RT scores have shown to be skewed [105], we used the median as an indicator of the central tendency [106]. As dependent variable for the analysis of the tACS effect on the spatial cueing task, we used the visuospatial attention bias score. For this, we subtracted the inverse efficiency score (RT/accuracy) [107,108] of right from left target location trials per condition.

#### Detection task

To investigate the effects of tACS on the detection task performance, we calculated two different bias scores. The contrast threshold bias score was calculated by subtracting the threshold for left from right targets. For the computation of the bias in indicated target location, we analysed the bilateral target trials in which an incorrect response was given. Here, we subtracted the number of trials in which the participant mistakenly indicated that the target appeared on the right side from the number of trials in which (s)he indicated that it appeared on the left side.

#### Statistical analysis

Mixed model analysis, an increasingly popular statistical approach [109–112], was used to analyse the EEG and behavioural data in SPSS. We used a compound symmetry covariance structure, which is suited for within-subject designs. As follow-up analysis on significant main effects of stimulation condition, we performed planned comparisons between the sham and the active stimulation conditions. Bonferroni correction was used throughout to correct for multiple comparisons.

#### EEG

First, we determined the IAF test-retest reliability by calculating the IAF for each session and running an intraclass correlation on the IAF estimates. A Pearson correlation analysis was used to analyse the association between age and the mean IAF over all sessions. Furthermore, we tested whether the IAF was shifted towards the stimulation frequency by fitting a mixed model on the IAF change score (IAF_post-measurement_ – IAF_pre-measurement_) with stimulation condition as factor. For the analysis of the neural stimulation effect, we fit a mixed effect model with stimulation condition as factor and ALI as dependent variable. As follow-up analysis on the full model, we conducted several pairwise comparisons. To find out which hemisphere drives the alpha power lateralization effect, we ran an additional analysis on the percentage increase in alpha power per hemifield using stimulation condition as factor.

#### Spatial cueing task

We first analysed the cueing effect in the spatial cueing task by fitting a mixed model on the median RT scores of the sham condition using Type of Cue as a factor. Then we analysed the tACS effect on the visuospatial attention bias score. The endogenous task was implemented as a 4 (stimulation condition: IAF, IAF+2Hz, IAF-2Hz, sham) x 3 (type of cue: valid, neutral, invalid) within-subject design. Mixed effect models were fitted on the visuospatial attention bias scores including stimulation condition and type of cue as factors and the electrophysiological entrainment effect ALI as a covariate. As follow-up analysis on significant interaction effects with the covariate, we conducted simple slope analyses to estimate the categorical condition effect at different levels of the continuous covariate [113,114]. To this end, we calculated three new covariate variables: alpha_mean,_ alpha_low_ and alpha_high_. To this end, we first determined the mean and standard deviation of the original covariate variable. Alpha_mean_ is the centred version of the original covariate and was computed by subtracting the precalculated mean from each individual score. Alpha_low_ and alpha_high_ were determined by adding or subtracting one precalculated standard deviation from each individual score of the centred covariate respectively [113,114]. Subsequently, three mixed effect models, one per new covariate, were fitted, using stimulation condition and type of cue as factor and visuospatial attention bias as dependent variable. Per stimulation condition, we omitted observations with a particularly low accuracy score (<55%) and excluded estimates based on an insufficient number of trials (<10) per condition. As a result, 1.2% and 2.8% of the total amount of observations were deleted respectively. Furthermore, we ran linear regression analyses per stimulation condition with ALI as predictor and visuospatial attention bias as dependent variable. As control analysis, we subsequently tested whether a model including age or the IAF as additional predictors is superior to a model with only ALI as predictor. To this end, we ran mixed model analyses on the visuospatial attention bias score with and without the additional predictors and compared the fit of the different models using log likelihood tests. One influential case with a cook’s distance above 1 was excluded.

#### Detection task

The detection task was implemented as a within subject design with one factor (stimulation condition: IAF, IAF+2Hz, IAF-2Hz, sham). We ran two mixed model analyses score using stimulation condition as factor and ALI as covariate. For the first analysis we used the threshold bias and for the second analysis the bias in indicated target location as dependent variable. Estimates based on an accuracy below 40% or above 60% were omitted to guarantee comparable contrast thresholds.

#### Blinding success

We administered a questionnaire at the end of each session, which prompted the participants to evaluate whether real or sham stimulation was applied. To statistically verify that blinding was maintained, we fitted generalized linear equations on the rated stimulation conditions using the actual stimulation condition (IAF, IAF+2Hz, IAF-2Hz, sham) as factor. The rated stimulation condition was assessed on an ordinal scale with seven levels ranging from ‘I definitely experienced placebo/sham stimulation’ to ‘I definitely experienced real stimulation’. A chi square analysis was used to test whether the actual stimulation condition affected the rated stimulation condition.

---

**Fig S1.**
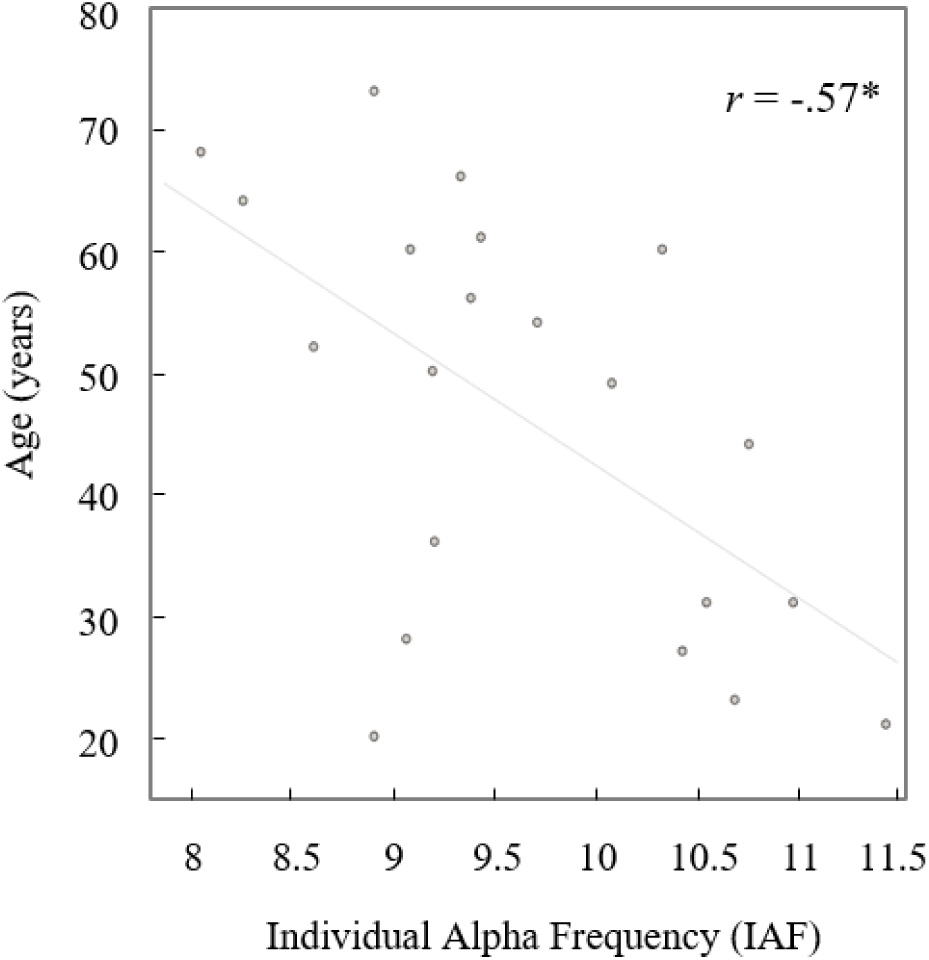
Association between IAF and age. Each point depicts the data of one participant. The asterisk marks a significant correlation with a p-value ˂ 0.05. The r-value indicates the correlation coefficient.

**Fig S2.**
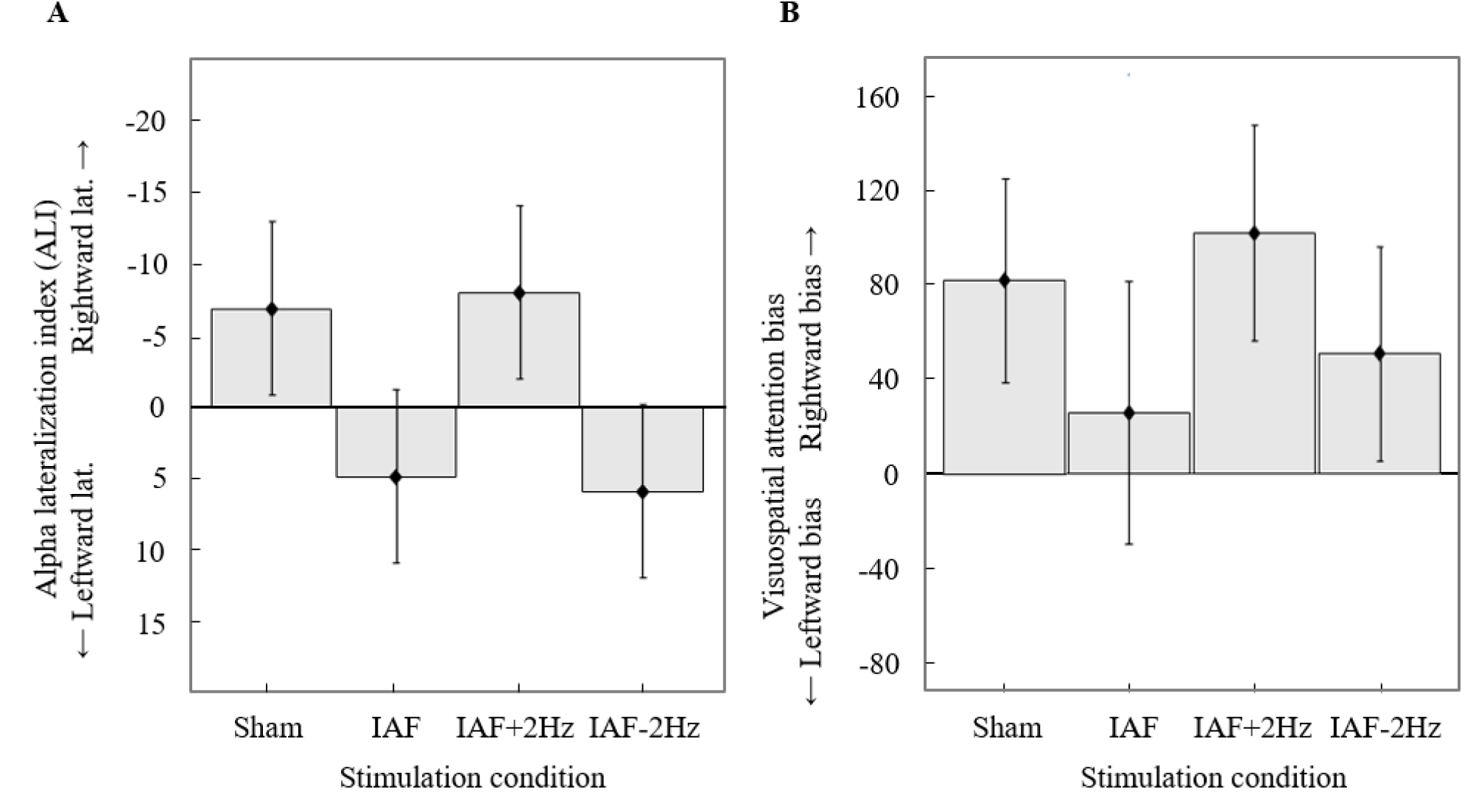
Electrophysiological and behavioral stimulation effect including all three minutes of the EEG post-measurement. (A) *Alpha lateralization index (ALI) per stimulation condition*. The main effect of stimulation condition (F(3,60) = 1.94, p = .132) and the pairwise comparisons (IAF tACS vs sham: t(60) = -1.56, p = .125; IAF+2Hz tACS vs sham: t(60) = .15, p = .881; IAF-2Hz tACS vs sham: t(60) = -1.70, p = .095) were not significant. (B) *Visuospatial attention bias in the spatial cueing task per stimulation condition and for a high value of the covariate ALI*. The interaction effect between stimulation condition and ALI (F(3,201.56) = 1.02, p = .384) and the pairwise comparison for a high value of ALI (IAF tACS vs sham: t(202.34) = 1.23, p = .221; IAF+2Hz tACS vs sham: t(199.23) = .66, p = .511; IAF-2Hz tACS vs sham: t(200.37) = .99, p = .322) were not significant. The error bars depict the pooled standard error of the mean across participants.

**Fig S3.**
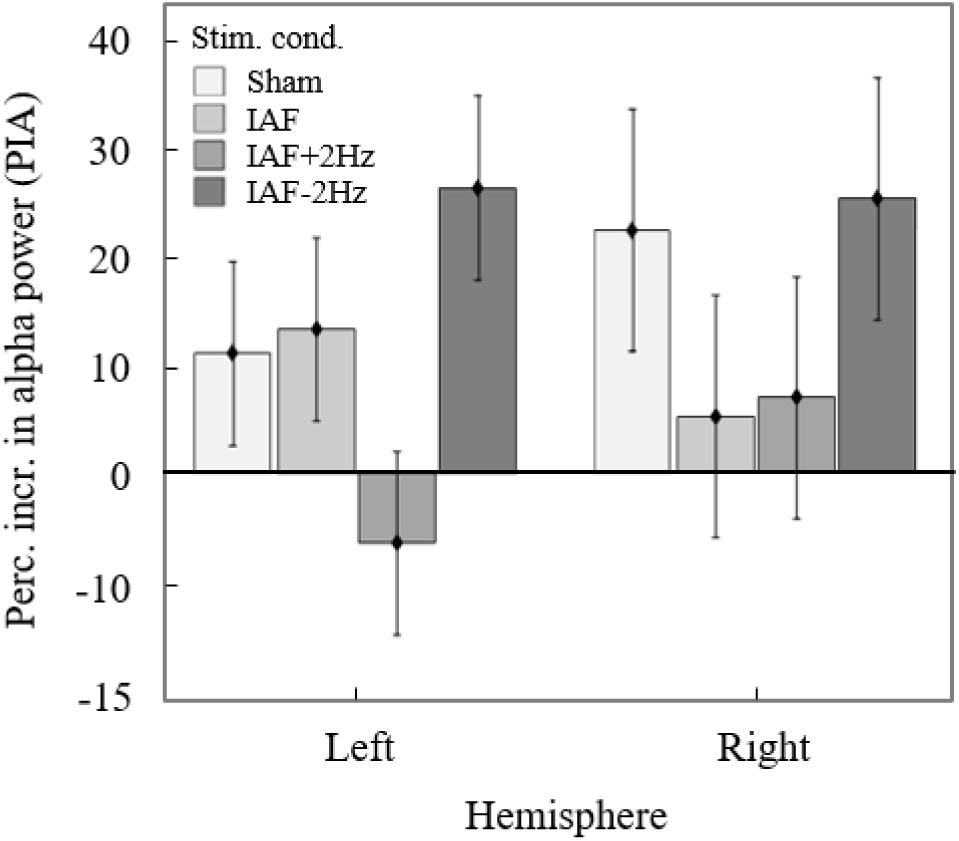
Percentage increase in alpha power (PIA) from the pre- to the post-measurement per hemisphere and stimulation condition. The data of the sham, IAF, IAF+2Hz, IAF-2Hz and sham stimulation conditions are depicted in increasingly darker shades of grey. The main effect of stimulation condition was significant for the left (F_3,60_ = 2.84, p = .045) but not for the right hemisphere (F_3,60_ = 1.22, p = .312). However, all planned comparisons for the left (IAF vs sham: t_60_ = .20, p = 1.00; IAF-2Hz vs sham: t_60_ = -1.54, p = .635; IAF+2Hz vs sham: t_60_ = 1.35, p = .505) as well as the right hemisphere (IAF vs sham: t_60_ = -1.30, p = 1.00; IAF+2Hz vs sham: t_60_ = -1.16, p = 1.00; IAF-2Hz vs sham: t_60_ = 0.22, p = 1.00) were not significant. Hence, we could not reveal whether the alpha power lateralization effect is driven by alpha power changes in the left or right hemisphere. The error bars depict the pooled standard error of the mean across participants.

**Fig S4.**
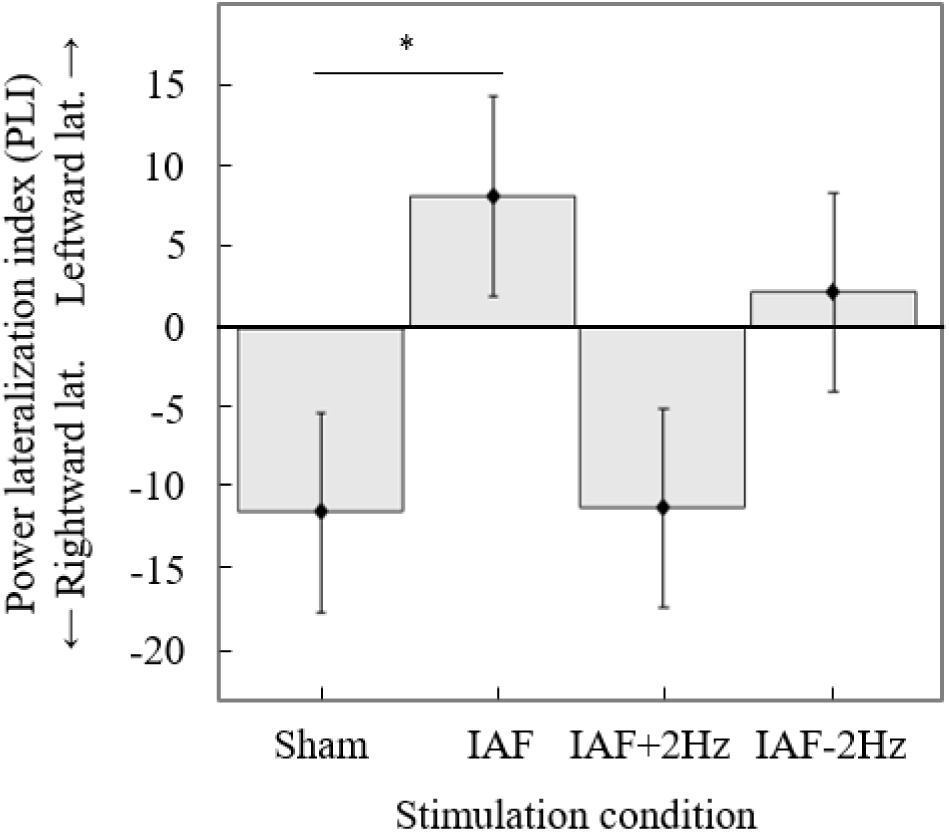
Power lateralization index (PLI) per stimulation condition. A positive value of PLI means a greater increase in power at and around the stimulation frequency in the left relative to the right hemisphere. The lines and asterisks indicate the results of planned comparisons between sham and the active stimulation conditions. The asterisks mark p-values < .05 and the error bars depict the pooled standard error of the mean across participants.

**Fig S5.**
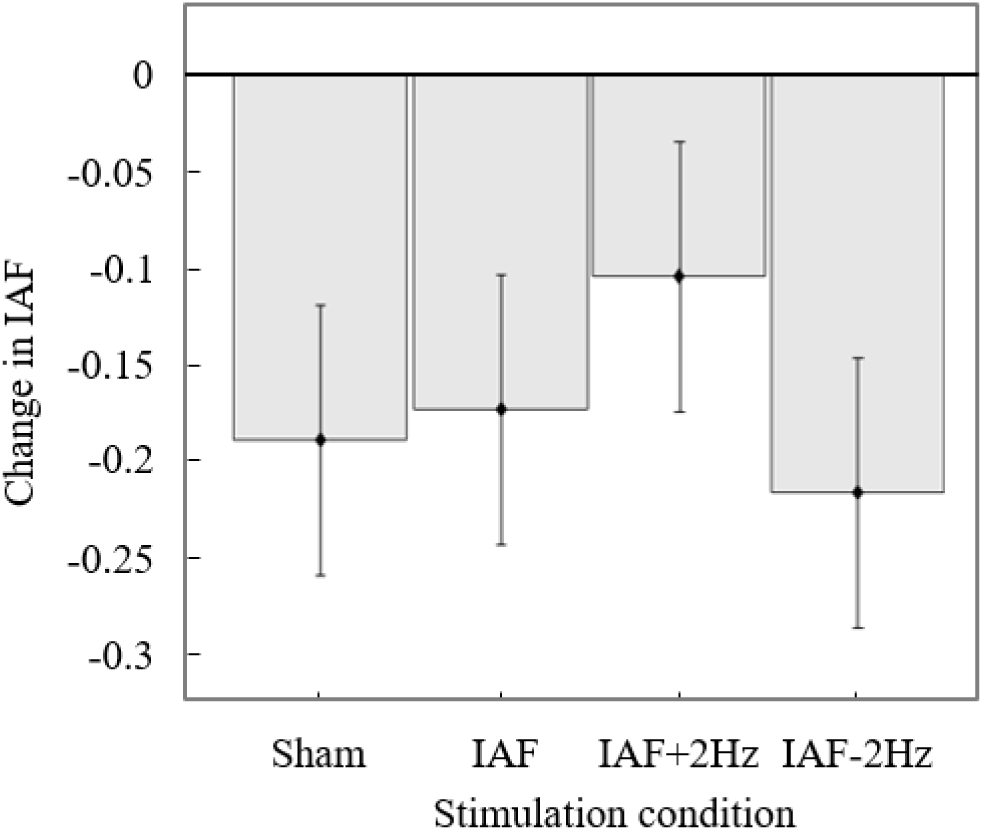
Change in IAF per stimulation condition. A negative value indicates a decrease in IAF from the pre-to the post-measurement. Error bars depict the standard error of the mean across participants.

**Fig S6.**
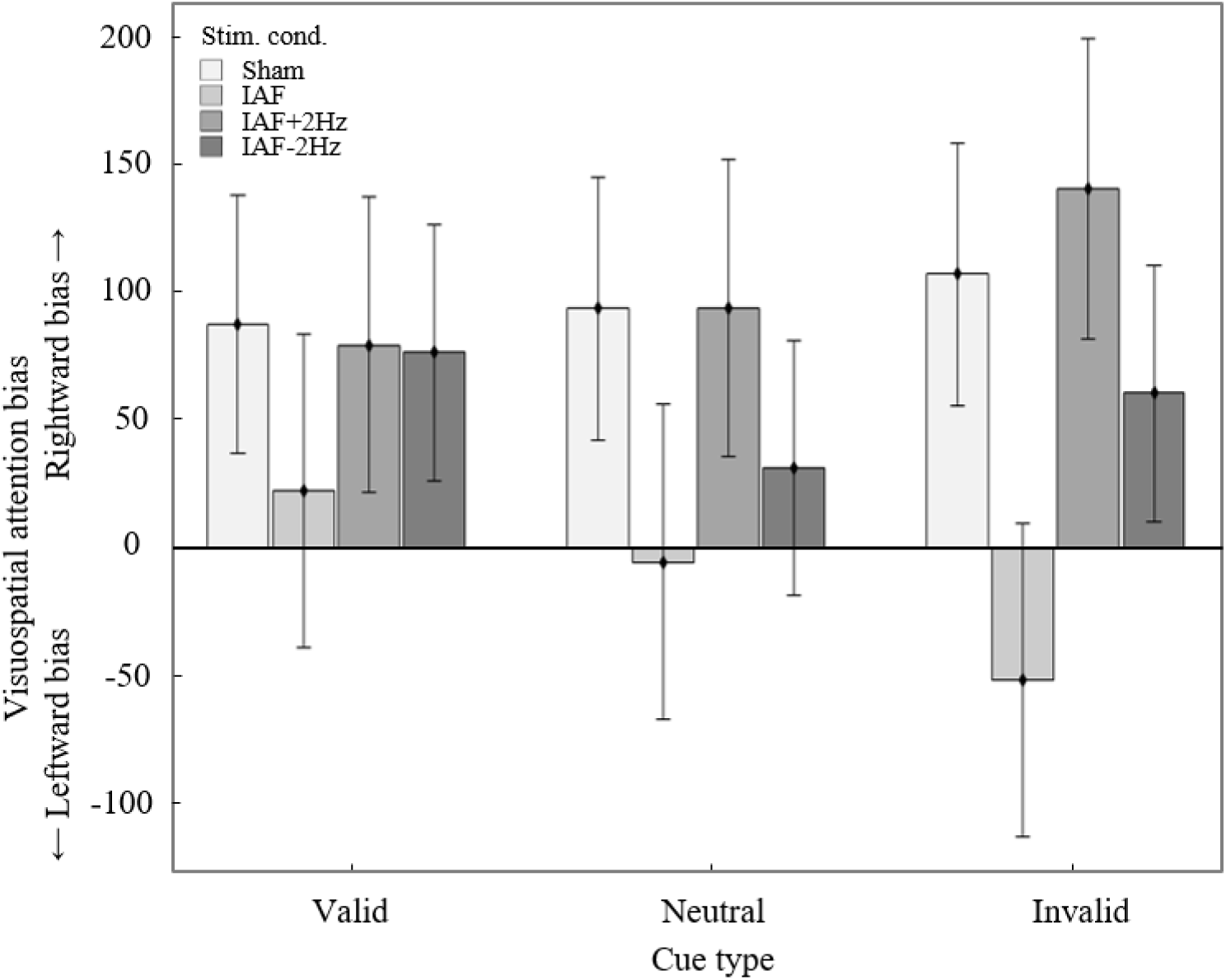
Spatial cueing task: Visuospatial attention bias per stimulation condition and cue type for a high value of the covariate ALI. A positive value of the visuospatial attention bias means that participants were less efficient (reaction time/accuracy) in responding to target stimuli in the left relative to stimuli in the right target location. The error bars depict the pooled standard error of the mean across participants.

**Fig S7.**
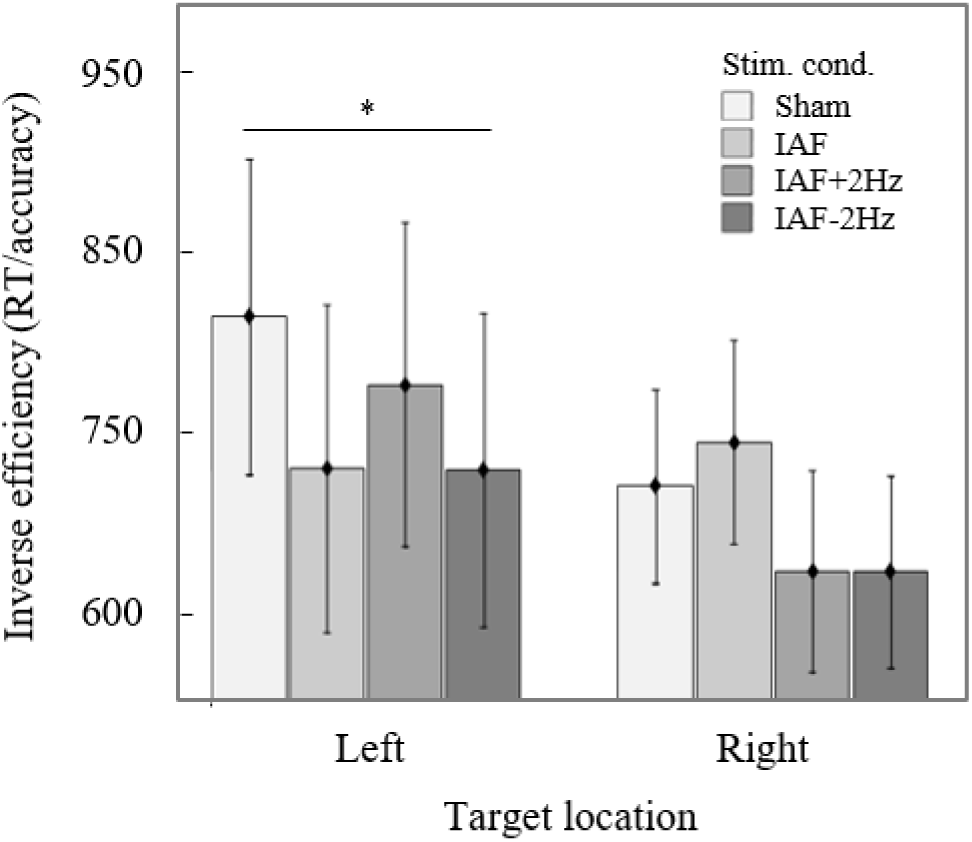
Spatial cueing task: Inverse efficiency (reaction time/accuracy) score per target location and stimulation condition for a high value of the covariate ALI. The data of the sham, IAF, IAF+2Hz, IAF-2Hz and sham stimulation conditions are depicted in increasingly darker shades of grey. There was a significant stimulation condition effect for the left (F_3,198.62_ = 2.94, p = .034) as well as the right target location trials (F_3, 199_ = 3.25, p = .023). For the left targets, the inverse efficiency score was lower for the IAF-2Hz condition (M = 728.70, SE = 86.83) as compared to sham (M = 814.33, SE = 87.29) (t_198.58_ = 2.75, p = .045). However, there were no other significant differences for the left (IAF vs sham: t_198.50_ = .03, p = 1.00; IAF+2Hz vs sham: t_198.11_ = .90, p = 1.00) or the right (IAF vs sham: t_198.68_ = .23, p = 1.00; IAF+2Hz vs sham: t_198.12_ = .65, p = 1.00; IAF-2Hz vs sham: t_198.34_ = 1.41, p = .805) target location trials. Asterisks marks p-values < .05 and the error bars depict the pooled standard error of the mean across participants.

**Fig S8.**
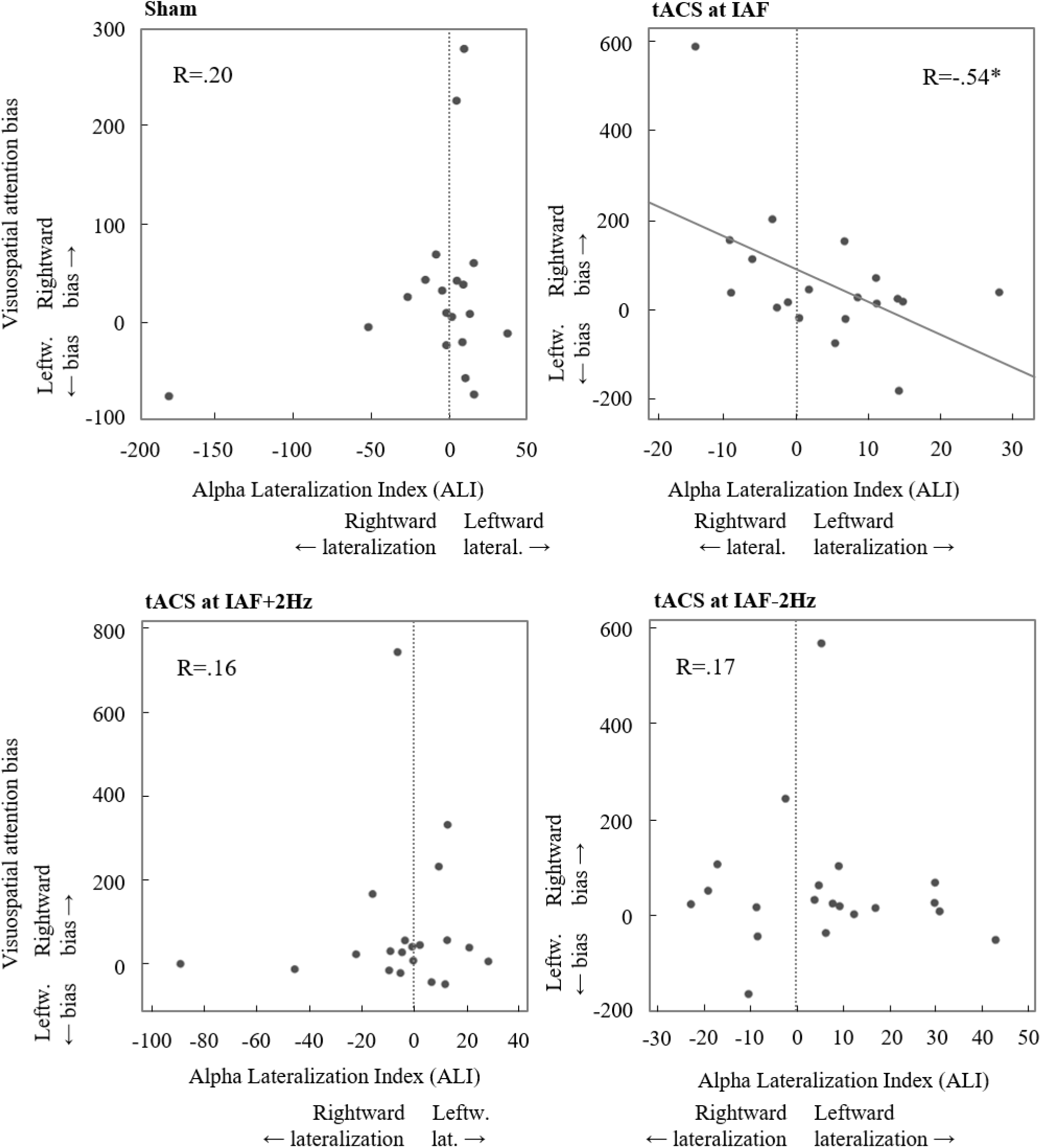
Association between ALI (including all three minutes of the EEG post-measurement) and visuospatial attention bias per stimulation condition. Per subplot, each point depicts the data of one participant. The alpha lateralization index (ALI) predicted the visuospatial attention bias in the IAF (*b* = -.54, p = .017) but not in the IAF+2Hz (*b* = .16, p = .521), IAF-2Hz (*b* = -.17, p = .502) or sham condition (*b* = .20, p = .392). The asterisk marks a significant effect with a p-value ≤ 0.05 and the vertical dashed line visualizes the zero value on the x-axis. The R-value indicates the regression coefficient.

**Fig S9.**
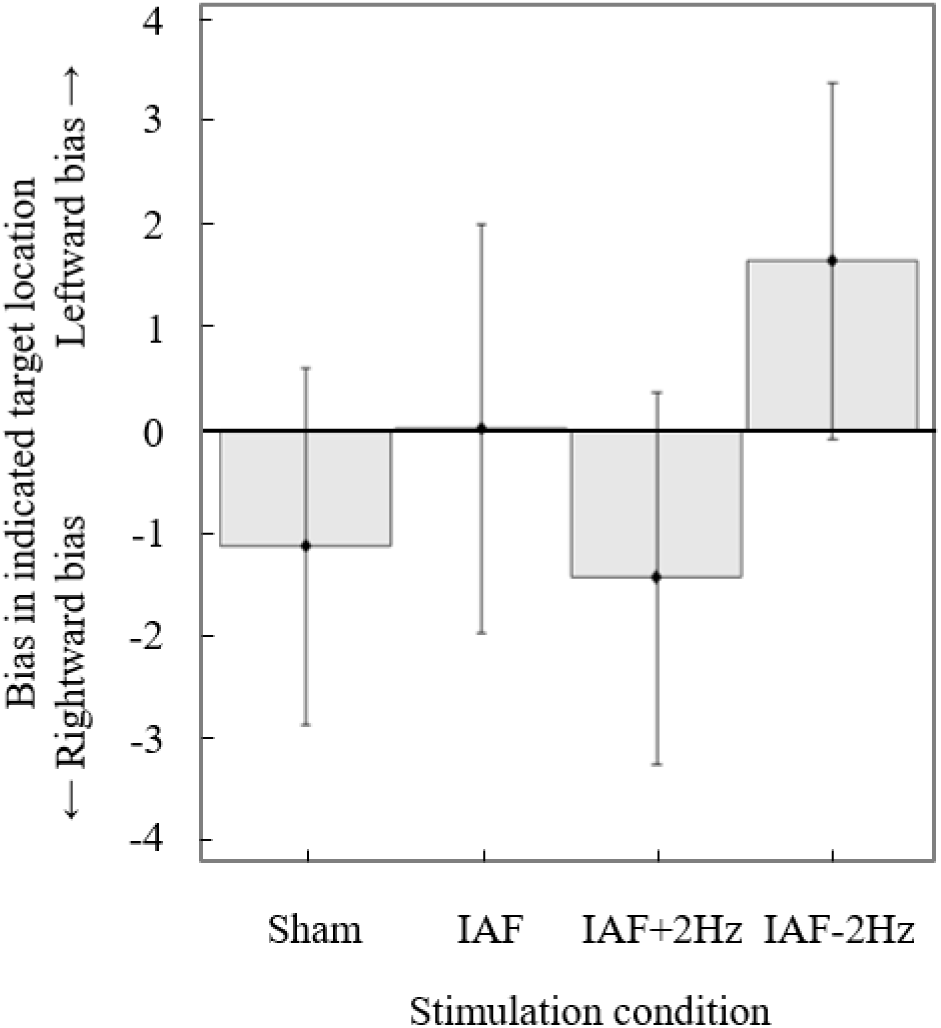
Detection task: Bias in indicated target location in bilateral target trials per stimulation condition. In some bilateral target trials, participants mistakenly responded that there was a unilateral stimulus in either the left or right hemifield. A positive bias score means that participants indicated more often that the target was in the left than in the right hemifield (leftward bias). The error bars depict the pooled standard error of the mean across participants.

